# Exponentially few RNA structures are designable

**DOI:** 10.1101/652313

**Authors:** Hua-Ting Yao, Mireille Regnier, Cedric Chauve, Yann Ponty

**Affiliations:** LIX, UMR 7161, Ecole Polytechnique Palaiseau, France; Department of Mathematics, Simon Fraser University Burnaby, Canada

**Keywords:** RNA Design, Inverse Folding, Pattern Matching, Analytic Combinatorics, Neutral Networks

## Abstract

The problem of RNA design attempts to construct RNA sequences that perform a predefined biological function, identified by several additional constraints. One of the foremost objective of RNA design is that the designed RNA sequence should adopt a predefined target secondary structure preferentially to any alternative structure, according to a given metrics and folding model. It was observed in several works that some secondary structures are undesignable, *i.e.* no RNA sequence can fold into the target structure while satisfying some criterion measuring how preferential this folding is compared to alternative conformations.

In this paper, we show that the proportion of designable secondary structures decreases exponentially with the size of the target secondary structure, for various popular combinations of energy models and design objectives. This exponential decay is, at least in part, due to the existence of undesignable motifs, which can be generically constructed, and jointly analyzed to yield asymptotic upper-bounds on the number of designable structures.

## 1 INTRODUCTION

RiboNucleic Acids (RNAs) are ubiquitous biomolecules equipped with a capacity of performing a wide variety of functions, both as a messenger enabling gene synthesis (mRNAs), as a regulator of gene expression (miRNAs…) or as a direct performer of a large collection of enzymatic activities (ncRNAs) [28]. For a large subset of ncRNA family, the adoption of a predefined structure is instrumental to the function(s) of individual molecules [45], and even, at times, the survival of its hosts organisms [20]. Accordingly, the evolutionary pressure on RNA families induced by RNA structure, at the secondary structure level, is at the core of most approaches for the identification of novel ncRNA families [46]. Improved characterizations of this pressure for individual represents a key challenge of RNA Bioinformatics and the object of current work, for instance in the case of the elusive long non-coding RNAs (lncRNAs) [38].

This strong connection between RNA structure and function has motivated the continuous development of mature computational methods for structure prediction [24, 34, 49]. More recently, researchers have attempted to harness the success of folding prediction approaches, and tackled a *de novo* design of structured RNAs. In its historic setting [24], the RNA Design, or inverse folding, consists in designing a sequence of nucleotides, folding into a predefined structure according to a criterion which can computed using available computational methods. RNA design is now an established problem in RNA bioinformatics, motivated by applications ranging from synthetic biology [42] to RNA therapeutics [47] through systems biology [14] and nanotechnologies [21].

Computationally, RNA design is a hard problem [4], motivating the development of several design methodologies [6] relying on exact exponential algorithms (constraint-programming [19], SAT solving) on heuristics (local search [1, 2, 5, 24, 48], genetic algorithms [13, 32], sampling [37]…). While the former methods are limited in their scope of applications by their extreme computational demands, methods of the latter category have encountered numerous applied successes [23], and enjoy a growing popularity. Historic objectives of design include the adoption, by the produced sequence, of a structure having energy as close as possible to the Minimal Free-Energy (MFE) achieved by the sequence. Modern formulations also include the minimization of defects, properties of the thermodynamic equilibrium that indicate a notion of distance to the expectation of a perfect design [9]. Those include the *probability defect* [9], the probability of not folding into the target structure, or the *ensemble defect* [48], the expected base pair distance to the target at the thermodynamic equilibrium.

However, some secondary structures do not admit a solution to the design problem. This was first observed by Aguirre-Hernández *et al* [1], with the discovery of two *undesignable* motifs, motifs for which alternatives would always be preferred by the usual Turner energy models [43]. This claim was later generalized to simple basepair based energy models by a study of a combinatorial version of RNA design [22], exhibiting motifs whose presence within a structure precludes its designability. However, the prevalence, in the folding space, of such undesignable motifs, and their impact on the overall combinatorics of designable structures, was never been assessed to date. Moreover, a characterization of undesignable structures would allow for sanity checks within design methods, avoiding the costly execution of a heuristics-based algorithm in contexts where no such solution exists.

Another motivation for this work pertains to theoretical evolutionary studies, where the RNA sequence to structure relationship represents an attractive model of neutral network [18], and a crucial conceptual framework to quantify the evolvability of species [10]. Indeed, the sequence/structure relationship in RNA enables the existence of, possibly large and highly diverse, subsets of sequences (genotype) folding into the same structure (phenotype), thus achieving the same fitness level. Studies of RNA neutral networks [17, 40] often require an enumeration of accessible phenotypes, *i.e.* the number of RNA secondary structures of a given size which are adopted as the most stable structure for *some* sequence. Since no exact method is known to compute this quantity, studies rely on available asymptotic estimates for the number of all secondary structures [44]. Such an implicit assumption of universal designability may bias studies [27, 29] of the underlying evolutionary dynamics, by artificially inflating the cardinality of structural ensembles. It is thus crucial to provide more precise (approximate) expressions for the number of designable structures.

In this work, we show that the existence of small undesignable motifs, which we call *local obstructions*, constitutes an intrinsic feature of RNA design objectives. An enumeration of the secondary structure that avoid those motifs thus represents an upper bound on the number of designable structures. A direct consequence of this observation is that the proportion of designable secondary structures is typically negligible beyond a certain sequence sizes. Indeed, a tree motif perspective on the problem, coupled with classic results in analytic combinatorics [16] imply that the proportion of designable structures over *n* nucleotides scales like *α*^*n*^, where *α* < 1 can be numerically computed from any collection of local obstructions. As a side product of our automated method for computing local obstructions, we are also able to establish a list of likely candidate sequences for each motifs of a given size.

After dedicating Section 2 to formal definitions for the key concepts, and state our main result, we describe in Section 3 the application of our general strategy on a simple combinatorial version of RNA design. We then show in Section 4 how small local obstructions can be computed, for any combination of defect, tolerance and energy models. Section 5 introduces a generic specification for enumerating secondary structures that avoid a collection of local obstructions, and describes a simple numerical procedure to derive asymptotic equivalents for the number of such structures. Section 6 presents the results of our analysis of different design objectives, using the realistic Turner energy model.

## 2 BACKGROUND AND RESULTS OVERVIEW

### RNA secondary structure

An RNA can be abstracted as a sequence *w* ∈ ∑*, ∑ := {A, C, G, U}, of nucleotides, having length *n* := |*w*|. For a sequence *w*, a secondary structure is a set *S* of base pairs *i*, *j*, *i* < *j* ∊ [1, *n*], representing the interaction of nucleotides at positions *i* and *j* through hydrogen bonds, such that:

1. Base pairs are pairwise non-crossing, *i.e.* ∄(*i*, *j*), (*k, l*) ∊ *S* such that *i* < *k* < *j* < *l*;
2. A minimal distance of *θ* is ensured between interacting positions, *i.e.* ∀(*i*, *j*) ∈ *S*, *j* − *i* > *θ*;
3. Any position of [1, *n*] is involved in at most one base pair.

Positions of [1, *n*] that are not involved in any base pair are called *unpaired*. In the following, we will denote by 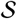 the entire set of secondary structures, and by _*n*_ its restriction to structures of length *n*.

A shown in Fig. 1, a secondary structure *S* of length *n* can be unambiguously represented as a rooted ordered tree *T* = (*V* := *V*_*i*_ ⋃ *V*_*j*_, *E*, whose nodes are either intervals [*i*, *j*] ∈ *V*_*i*_, *i* < *j*, representing base paired positions (*i*, *j*) in *S*, or singletons {*i*} ∊ *V*_*j*_, representing an unpaired position *i* in *S*. Note that leaves of such tree represent only singletons. Any edge (*u* → *v*) ∈ *E* connects intervals such that *u* ⊂ *v* and ∄*v*′ ∈ *V*_*i*_ such that *u* ⊂ *v*′ ⊂ *v*.

**Figure 1:**
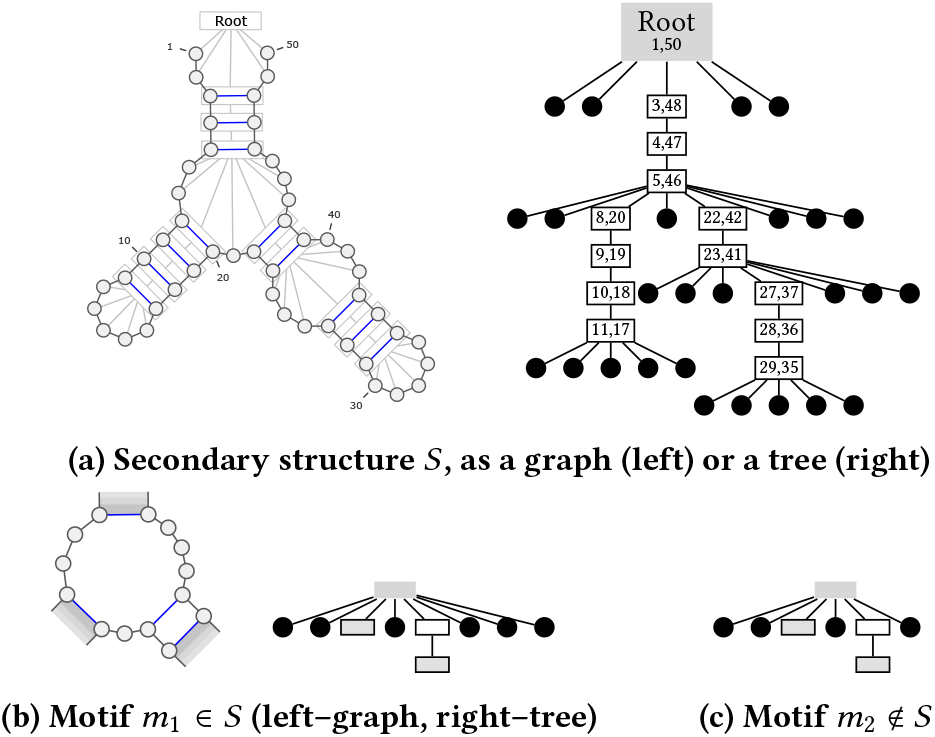
Graph and tree representations of an RNA secondary structure *S* of size 50 (1a). It features an occurrence of the motif *m*_1_ (1b), having size 14 with 2 paired leaves, rooted at node (5, 46). Although *m*_2_ (1c) resembles *m*_1_, it lacks two unpaired nodes and is thus does not strictly occurs at position (5, 46).

### Energy model

An energy model assigns a free-energy value to each pair (*w*, *S*), where *w* is an RNA sequence and *S* is a secondary structure for *w*. Popular energy models for RNA folding prediction, such as the base pairs-based *Nussinov* and the *Turner* nearest-neighbors models, consider additive contributions associated with the *shallow subtrees*, *i.e.* subtrees of depth 1 occurring in *S*, and their respective nucleotides assignments. Hence, an *energy model* is a function 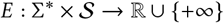 such that

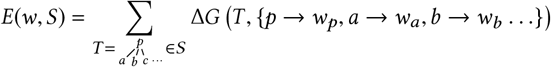

where Δ*G* (*T*, *m*) is the free-energy, expressed in kcal.mol^−1^ associated with the assignment *m* of concrete nucleotides from *w* to the (pairs of) positions in the subtree *T*. In practice, values taken by Δ*G* are tabulated or extrapolated from experimentally-measured values.

### RNA Folding

RNA structure modeling aims, given a sequence *w*, to find one or several folding(s) of *w* into RNA secondary structure(s). Several paradigms exist, associated to different objective functions measuring the quality of a folding. In the energy minimization setting, a central algorithmic question is to compute the *minimum free energy* (MFE) structure(s), optimizing the thermodynamic stability:

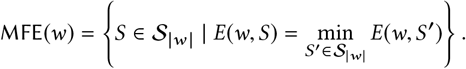

A second, increasingly popular, paradigm strives to predict structures that are representative of the Boltzmann ensemble of low energy structures. Under the hypothesis of a Boltzmann equilibrium, statistical mechanics postulates that, for a given sequence *w*, the putative secondary structures follow a Boltzmann distribution

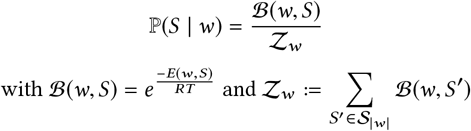

where *R* is the Boltzmann constant, *T* is the temperature,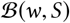 is called the *Boltzmann factor* of *w* and *S*, and 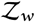 the *partition function* of *w*. Similarly, the probability of a base pair is defined as

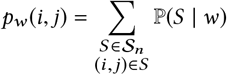

and *p*(*i*, *i*) represents the probability of *i* being left unpaired.

Note that, while an MFE structure has maximum probability in the Boltzmann ensemble, its probability can be arbitrary low, so achieving a high probability is not reducible to being an MFE. In fact, modern approaches typically elect structures that are, on average, maximally similar (MEA [31], centroids [8]) to random structures in the Boltzmann ensemble.

### Defects and negative RNA design

Given a *target secondary structure S*^*^, the *negative RNA design* problem, also called inverse folding, consists in producing one or several RNA sequences *w* that folds into *S*^*^ while avoiding alternative folds of similar quality for the chosen energy model.

The avoidance of alternative structures is captured by a notion of *defect*, defined as a function 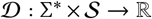. RNA design methods usually consider one of the three following defects:

1. The *Suboptimal Defect* 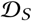 of a sequence *w* is defined as the energy distance to the first suboptimal, such that

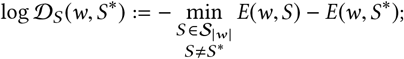
2. The *Probability Defect* 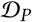 represents the probability of folding into any other structure than *S*^*^:

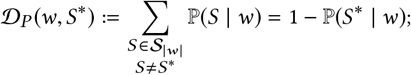
3. The *Ensemble Defect* 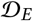 is the expected base pair distance between *S*^*^ and a random structure, generated with respect to the Boltzmann probability distribution:

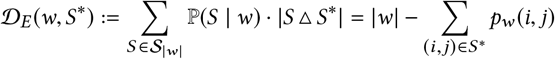

with |*S Δ S′*| a shorthand for the set symmetric distance, also known as base pair distance.

Now we can define the *main objectives of negative RNA design*. Given a real-valued threshold *ε* ≥ 0 and a defect 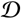, a sequence *w* is a *(negative)* 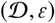-*design* for a structure *S*^*^ if and only if

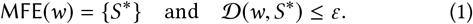

Similarly, we call 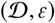-designable a secondary structure that does admit at least a valid design. Note that the defect definition also depends on the chosen energy model, but we chose to make this dependency implicit for the sake of simplicity.

### Motifs and local defect

A *motif* is a rooted ordered tree, similar to a secondary structure, but whose leaves may represent base paired positions. We say *a motif m occurs in a secondary structure S (resp. a motif m*′) or *a secondary structure S (resp. a motif m*′) *contains a motif m* if *m* is a subtree of *S* (resp. *m*′), rooted at any base paired node in *S* (resp. *m*′) and obtained by deleting all the children for a subset of its base paired nodes. In other words, a node in *m* either has exactly all of its children within *S* (resp. *m*′), or none. See Fig. 1 for an example.

Consider a motif *m*, having a root base-pair (*i*, *j*) and paired leaves (*i*_1_, *j*_1_), …, (*i*_l_, *j*_l_), and let *w*, |*w*| = *n*, be an assignment of nucleotides to the positions of *m*. We define the *local defect* 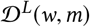 similarly as 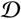, by replacing 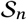 with

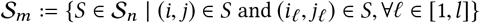

a restricted set of structures where both the root, and all the paired leaves, of *m* appear as base pairs. A crucial observation, of which we omit a formal proof in the interest of space, is stated in the following proposition.

#### PROPOSITION 1.

*For any defect* 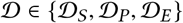, *sequence w*, |*w*| = *n, and structure*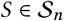, *one has*

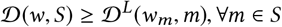

*where w*_*m*_ *is the restriction of w to the positions in m*.

#### COROLLARY 1.

*If there exists a motif m* ∈ *S*^*^ *such that*

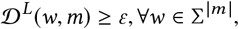

*then S*^*^ *cannot be* 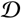*-designed*.

In other words, the presence in the target structure *S*^*^ of a motif that cannot be designed *locally* is sufficient to forbid the existence of a sequence *w* that would constitute a design for *S*^*^.

### Problem statement and results overview

In this work, we address the following question: *Given an energy model, a design criterion, how many secondary structures of a given length actually admit a negative design?* Our main result is summarized by the following theorem.

#### THEOREM 1.

*For any energy model, defect*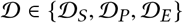*and tolerance ε* ≥ 0, *only an exponentially small fraction of the secondary structures in* 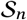 *are* 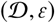-*designable*.

## 3 BASIC COMBINATORIAL DESIGN

Here, we consider the special case of the 0-dominance criterion in the simplest energy model, the Nussinov model, considered in a previous work [22]. In this setting, the design problem simplifies into finding an RNA sequence admitting a unique folding maximizing the number of base pairs, such that this folding coincides with the given target secondary structure.

### THEOREM 2.

*Let d*_*n*_ *be the number of secondary structures that are designable in the Nussinov model with θ* = 1. *Then*

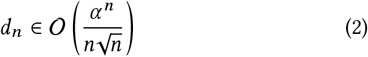

*where α* = 2.35 … *is the smallest positive real root of*

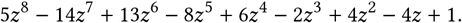

### COROLLARY 2.

*The probability that a uniform random secondary structure of length n, with θ* = 1*, is designable in the Nussinov model for the* 0*-dominance criterion is in* 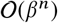, *where* 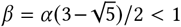.

To prove Theorem 2, we rely on analytic combinatorics techniques, widely used in analysis of algorithms [15] and bioinformatics [41], exposed in [16]. Their application in this context involves the following steps:

1. Identify a collection of secondary structure motifs 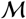 whose occurrence in *S* implies that *S* is not designable in the Nussinov model for the 0-dominance criterion;
2. Design a grammar for the set 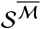 of all RNA secondary structures excluding this motif;
3. Derive and solve a system of functional equations satisfied by the generating function 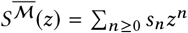, where *s*_*n*_ is the number of structures of 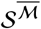 of length *n*;
4. Use singularity analysis to obtain an asymptotic equivalent for *s*_*n*_, in particular the coefficient *α* of (2) in Theorem 2, called the *growth factor*, that drives the exponential growth of *s*_*n*_ as a function of *n*.

Corollary 2 follows from Theorem 2 and the fact that, when *θ* = 1, the asymptotic number *t*_*n*_ of secondary structures of length *n* is such that

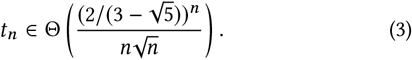

The exponential decrease in the proportion of designable secondary structures follows from 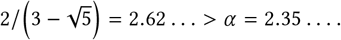.

We now turn to the proof of Theorem 2. For step (1) of the approach outlined above, we rely on the recent paper [22], where it was proved that a secondary structure cannot be designed if its tree representation includes an internal node whose children set contain ≥ 2 internal nodes and at least one leaf (the collection of motifs 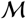 discussed above).

The set of tree representations of the secondary structure avoiding this local motif can be generated by the context-free grammar given below

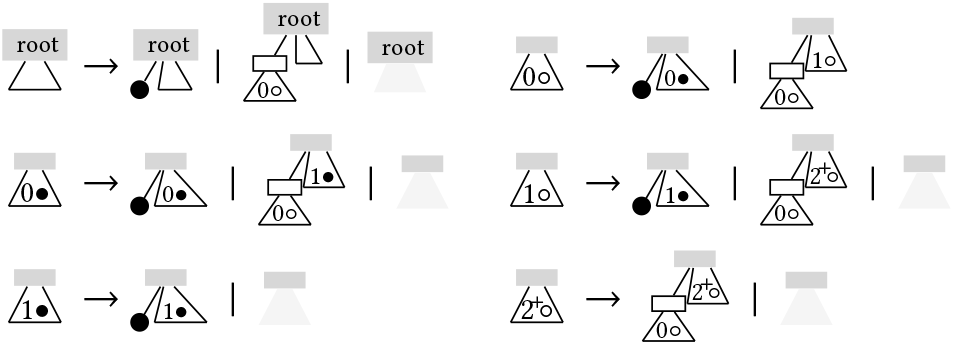

Intuitively, this grammar keeps track of properties of the structural elements generated for the current internal node. Except for the root, each non-terminal is indexed by pairs taken from (*i*, *u*) ∈ {0, 1, 2^+^} × {∘, •}, where, within the current siblings, *i* represents the number of internal nodes/base pairs, and *u* expresses whether (•) or not (∘) a leaf has been generated. Notice that the grammar implicitly excludes structures having a three siblings composed of two internal nodes and one leaf ((*i*, *u*) = (2^+^, •)), i.e. the motif 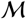.

Following standard enumerative combinatorics techniques that links combinatorial specifications to the calculus of generating functions [16], one obtains that the ordinary generating function 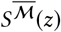 is defined by the system of functional equations below.

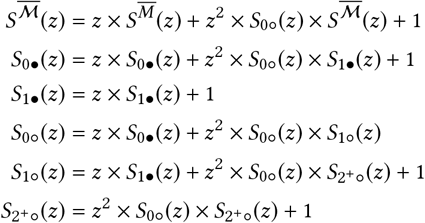

Solving the system using algebraic elimination, followed by a careful choice of the right conjugate, one obtains a closed form for the generating function 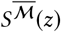:

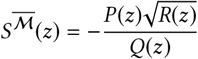

where

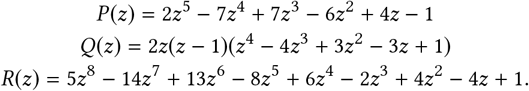

It follows from classic transfer theorems [15] that the singularity of this generating function is of square-root type, leading to the following asymptotic expansion for its coefficients,

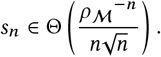

where 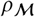 is the dominant singularity of *S*_r_(*z*), *i.e.* the smallest root of *R*(*z*), and can be numerically evaluated at 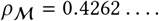

Theorem 2 follows from 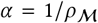 and the fact that *d*_*n*_ ≤ *s*_*n*_. Indeed, while it is necessary for a designable secondary structures to avoid 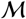, this condition is not sufficient.

## 4 LOCAL OBSTRUCTIONS

In this section, we describe an algorithm to compute local obstruction, motifs whose presence within a secondary structure forbids its design with respect to some predefined design objectives. Fig. 3 describes the main workflow of this study.

### 4.1 Emulating a local defect with constraints

The *minimal completion* of a motif *m* for a nucleotide assignment *w* is a pair (*S*_*m*_, *w*_*m*_) such that:

- *S*_*m*_ is the secondary structure obtained from *m* by adding *θ* unpaired nodes (leaves) under each paired leaf node;
- *w*_*m*_ is the sequence obtained by inserting *θ* occurrence of the letter A under paired leaves.

In this study, we set *θ* = 1 for the *Nussinov* model and *θ* = 3 for the *Turner* model. In other words, in the *Turner* model, the minimal completion is obtained by replacing any paired leaf ▭ by 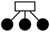. Figure 2 illustrates the application of the minimal completion to the motif *m*_1_ in Fig. 1b. A *trimming* operation is defined as the inverse of the completion, and allows to recover a motif/sequence pair from its completion.

**Figure 2:**
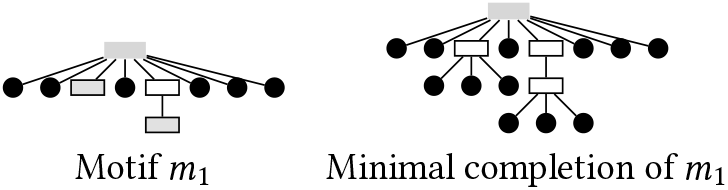
Minimal completion of the motif *m*_1_ (Fig. 1b).

**Figure 3:**
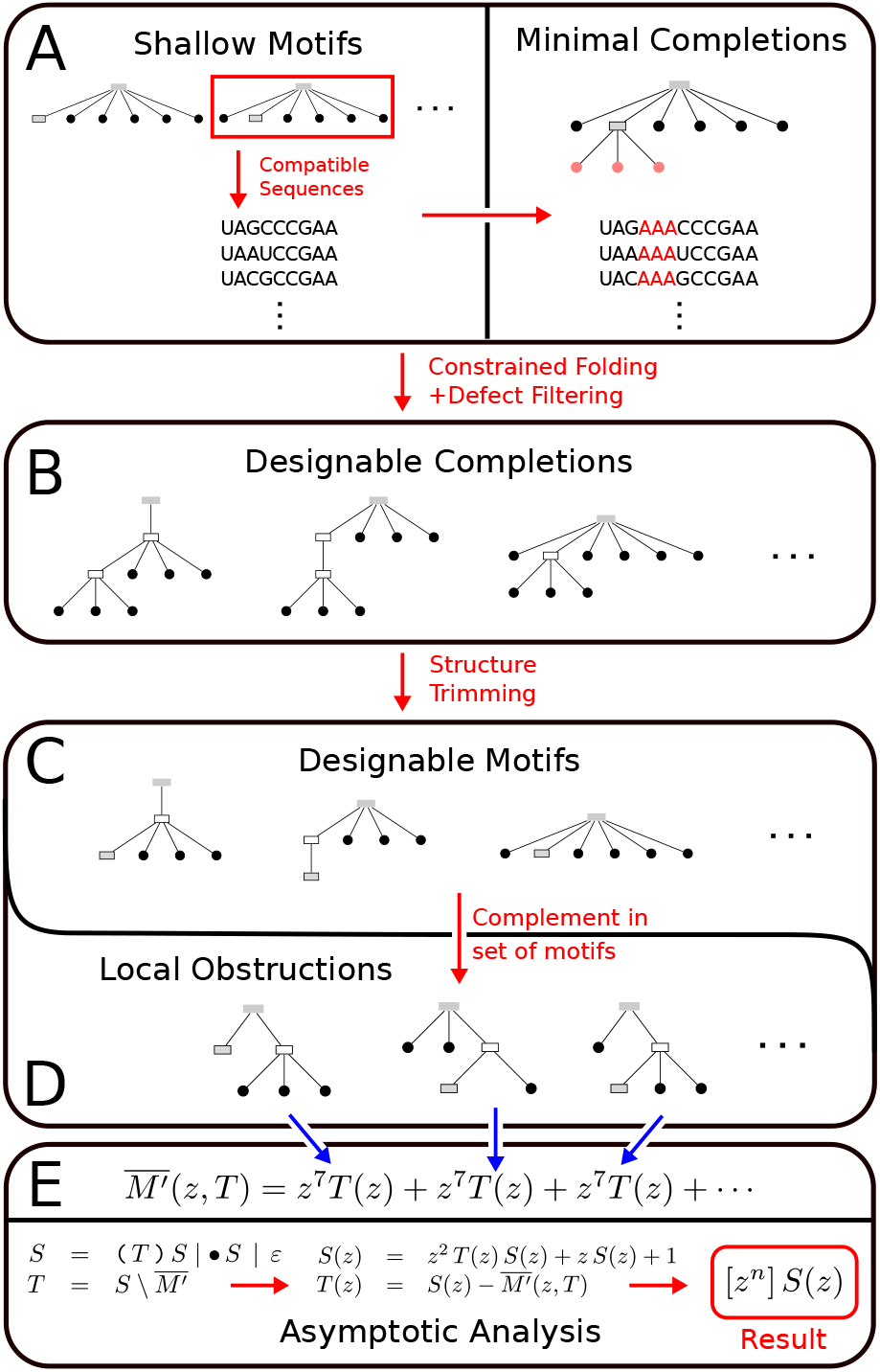
General workflow. Minimal completions are computed for all pairs of shallow motifs and compatible sequences (A). Constrained MFE predictions, filtered by defect, produce designs for refinements of the input motif (B). Designable completions are then trimmed into a set of designable motifs (C), itself complemented to obtain local obstructions (D). A grammar for secondary structures avoiding obstructions is built, and singularity analysis gives an asymptotic upper bound for designable structures (E).

Given a length *k*, a *folding constraint C* is a set consisting of positions from [1, *k*] and pairs from [1, *k*] ^2^, respectively representing positions forced to remain unpaired and paired to a specific partner. The *constrained defect* 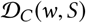 can be defined by restricting the computation to structures compatible with the constraint *C*. Such constraints are supported by reference implementations of energy-minimization and partition-function algorithms, and can be easily enforced in simpler energy models.

Consider a motif *m*, and its minimal completion (*S*_*m*_, *w*_*m*_), we define the *induced constraint C*_*m*_ of *m* as consisting of:

- the root base pair of *S*_*m*_;
- the base pairs in *S*_*m*_ stemming from the paired leaves in *m*;
- the unpaired positions introduced by the completion.

Intuitively, such a constraint will limit the alternative conformations, considered by the defect computation, to be consistent with the boundaries of the initial motif.

#### PROPOSITION 2.

*For every defect* 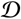 *and energy model considered in this work, one has* 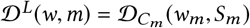.

In other words, the local defect of a motif can be practically computed by executing a constrained version of a, suitably constrained, global off-the-shelf algorithm (energy-minimization for 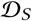, base-pair probability for 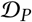 and 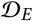) on the minimal completion of the motif. In particular, motifs that represent local obstructions to design, associated with large local defect, can be detected using this property as shown below.

### 4.2 Computing local obstructions

We now turn to the computation of a list of *local obstructions* over *k* nucleotides, motifs whose presence within any secondary structure implies that the overall defect exceeds a predefined *tolerance ε* ≥ 0.

In principle, we could compute all possible motifs and nucleotides assignments, followed by an evaluation of the local defect, as described in the previous section. We could then simply consider as local obstructions any motif which, for all its compatible sequences, fails to satisfy the *ε* defect threshold. Indeed, Prop. 1 implies that any motif whose local defect exceeds *ε* cannot be part of a secondary structure having defect less than *ε*, thus representing a local obstruction. Since motifs are essentially structures over *k* nucleotides (for *θ* = 0), the complexity of this approach would be in 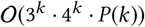, for *P*(*n*) the complexity of the (constrained) direct MFE/defect algorithm.

This complexity can be further reduced by restricting the above computation to *shallow motifs*, motifs having tree height 1 consistng of paired and unpaired nodes underneath a root node. For any sequence assignment to such a motif, running a constrained energy minimization algorithm on the minimal completion of the sequence either returns one, or multiple co-optimal solutions. In the case of multiple solutions, the sequence admits several alternative local MFE folds. It is thus not suitable for any *refinement* of the shallow motif, *i.e.* any motif that includes the pairs of the shallow motif. Conversely, a unique MFE solution is provably a refinement of the shallow motif, and one concludes that the motif is designable. Since every motif over *k* nucleotides is a refinement of some shallow motif, this strategy produces the same output as the above-described one. Its complexity, however, is reduced to 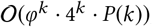, where 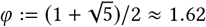 is the golden ratio, observing that shallow motifs are counted by the Fibonacci numbers.

Given a defect 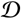 restricted to a value *ε*, and a given motif size *k*, our algorithm execute the following steps:

- Enumerate all *shallow motifs* (of depth 1) of size *k*;
- For any such motif *m*^◦^, consider any assignment *w*^◦^ consistent with the paired nodes in *m*^◦^:

− Build the minimal completion 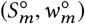 of (*m*^◦^, *w*^◦^), and execute on 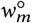 a constrained MFE folding algorithm, using the induced constraint 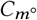;
− If the MFE computation returns a unique structure *m*⭑, consider the motif *m*′ obtained by trimming *m*⭑ (*m*′ is a refinement of *m*^◦^);
− Evaluate the local defect and, if 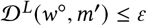, add *m*′ to the list 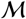 of designable motifs;
− Return 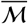, the set of all motifs of size *k* not in 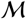.

A detailed version of the procedure is described in Alg. 1 and illustrated in Fig. 3.

**Algorithm 1:**
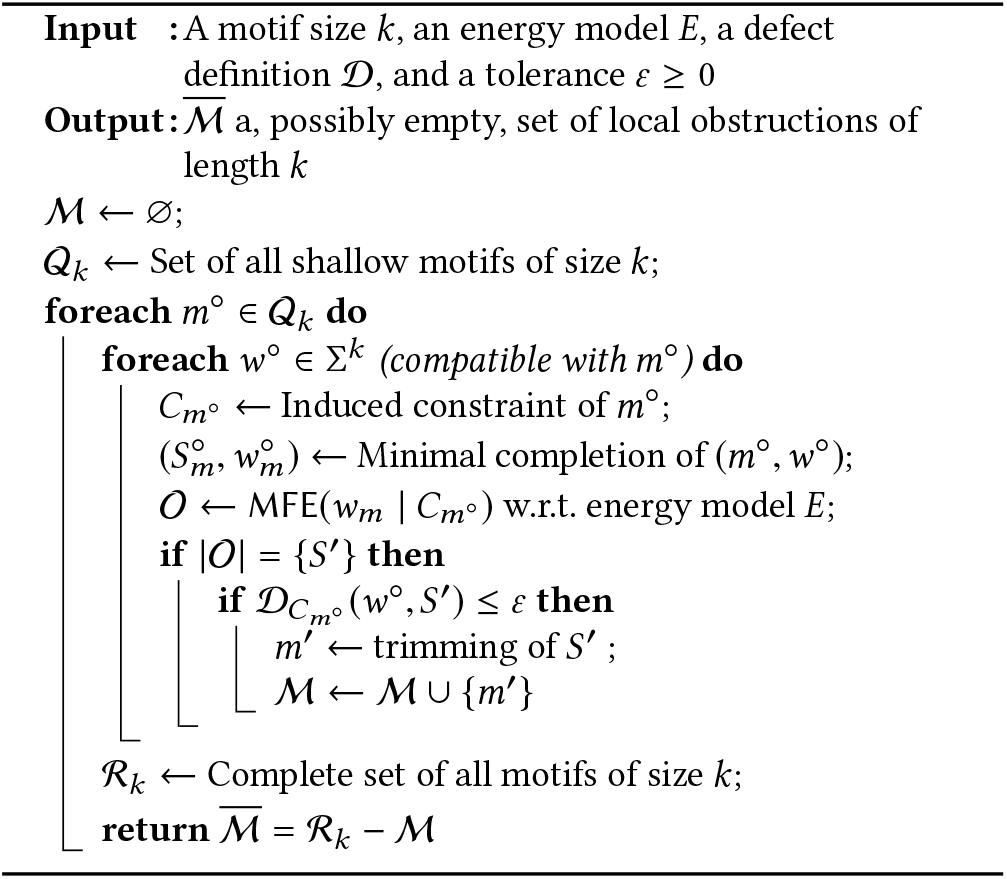
Computing local obstructions of a given size

#### PROPOSITION 3.

*Motifs returned by Alg. 1 are local obstructions*

PROOF. Fist, let us consider the properties of a motif *m* returned by the algorithm. Note that there exists only a single shallow motif *m*^◦^, of which *m* is a refinement. Since 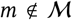 then, for each sequence *w*^◦^, either a lower constrained MFE fold was found, or the local defect exceeded *ε*. In the latter case, Proposition 1 implies that any pair (*S*, *w*), where *S* features *m*, and sequence *w* having nucleotide assignment *w*^◦^ on the motif positions, has defect greater than *ε*, thus *w* is not a design for *S*. In the former case where an alternative motif *m*′ is preferred to (or equally stable as) *m* for *w*^◦^, then for any structure *S* containing *m* and sequence *w*, having nucleotide assignment *w*^◦^ on the positions of *m*, a competitor to *S* for *w* can be constructed by replacing *m* by *m*′ in *S*. One concludes that, if 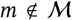, any structure *S*, *m* ∈ *S*, and sequence *w* does not represent a 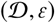-design.

The exhaustivity, for a given size *k*, of the list of motifs produced by Algorithm 1 remains unclear. Indeed, a motif is disregarded as a local obstruction as soon as its minimal completion folds correctly (with admissible defect) under suitable constraints for some sequence. Thus, there is no formal guarantee that a sequence would adopt this motif with an acceptable defect in the absence of constraints. However, we empirical observed that motifs not returned by the algorithm can overwhelmingly be included in design and, in particular, that the sequence of their minimal completion is a design for the completed structure. Moreover, the possible omission of some local obstructions is not overly critical, since our main goal is to provide upper bounds on the number of designable structures.

## 5 ENUMERATING SECONDARY STRUCTURES WHILE AVOIDING LOCAL OBSTRUCTIONS

Next, we turn to the computation of asymptotic equivalent for the number of secondary structures that avoid a collection of local obstructions, computed using the algorithm outlined in the previous section. We first start by eliminating redundant motifs, *i.e.* motifs that merely extend another motif, as shown in Figure 4, which can be done by running a classic tree alignment algorithm [26] in a pairwise fashion.

**Figure 4:**
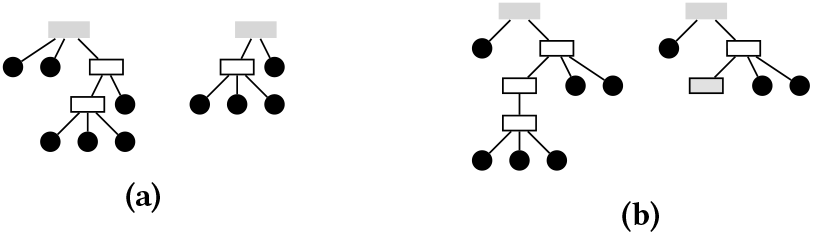
Two examples of pairs of redundant motifs. In both cases, the set of secondary structures rooted on the right motif strictly includes that of the right one, and we discard the left one from our computations.

### 5.1 Specification and generating function

Our approach represents an instance of the symbolic method [16], and is similar in essence to the detailed example of Section 3.

We establish that the set of all secondary structures avoiding a set 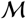 of local obstructions is generated by the following specification:

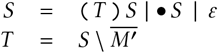

The first line essentially builds the set of all secondary structures (*θ* = 0), and is highly reminiscent of Waterman’s seminal decomposition [44]. The second line, however, subtracts the contributions of secondary structures which, when completed with a root, feature an occurrence of a local obstruction. In other words, *M*′ denotes the set of enclosed forests built from the inner part of local obstructions

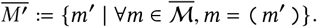

Therefore, the system of generating functions can be written as

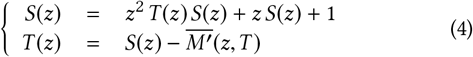

where 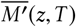 denotes the Ordinary Generating Function (O.G.F.) of the set of enclosed structure of motifs, defined as

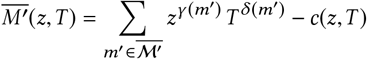

where *γ*(*m*′)(resp. *δ*(*m*′)) is the size (resp. number of paired leaves) of the motif *m*′, and *c*(*z*, *T*) is a correcting term to account for potential overlaps.

Indeed, in rare cases, some secondary structures may be counted in the O.G.F. associated to two or more motifs. Such structures would therefore be subtracted several times by the grammar, leading to an error while computing the singularity. Therefore, a correcting term *c*(*z, T*) is introduced, as described in Figure 5 to counterbalance the overcounting in such (rare) situations. Given the scarcity of such situations, we computed those terms manually for each pair of motifs. A more systematic solution could be implemented, using ideas from Collet *et al* [7], but the lack of immediate needs led us to leave this for an extended version of this extended abstract.

**Figure 5:**
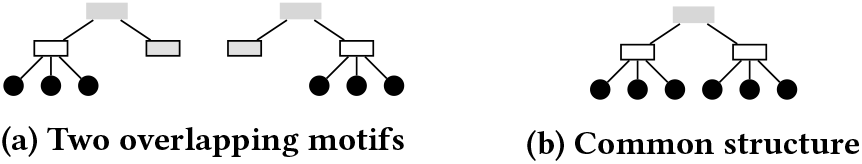
Both motifs in (5a) represent local obstructions, each of them contributing *z*^7^ *T* to the O.G.F.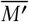(*z, T*). However, those motifs are overlapping, and any structure in the intersection, such as (5b), will be subtracted twice. We work around this issue by including a correcting term *z*^10^ in *c*(*z, T*).

**Figure 6:**
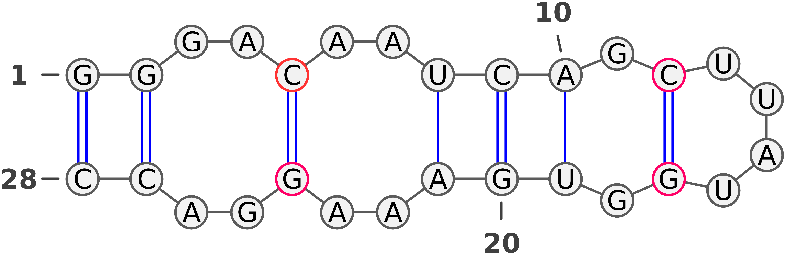
Example of secondary structure with isolated base pairs (red) in its MFE structure, obtained by connecting local solutions

### 5.2 Computing the dominant singularity

In general, one could use a symbolic calculus software (`Maple`) to solve the system (4), using some specialized package (`gfun` [39]) to extract the dominant singularity. However, in our case, such a approach turns out to scale poorly with the number of motifs and, more critically, paired leaves in the motifs (*i.e.* the degree of *T*(*z*) in 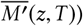). Therefore, we consider an alternative approach which combines an elementary symbolic calculus with a numerical determination of the dominant singularity.

Indeed, rewriting the system shows that *T*(*z*) is a solution of *G*(*z*, *y*) = 0 where

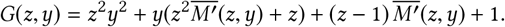

As 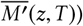 depends of 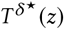, for 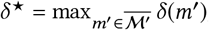, the degree of *G* with respect to *y* might be greater than 2, and the problem is not directly amenable to the techniques developed in Section 3. Nevertheless, it follows from this smooth implicit-function schema that *T*(*z*) is analytic. It is aperiodic and its dominant singularity, denoted *ρ*, is a non-zero root of *R*(*z*) defined as the resultant of two polynomials in *y*, namely:

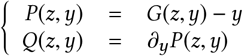

The solution is easily derived by a numeric approach. The generating function *S*(*z*) shares the same dominant singularity as *T*(*z*). Thus, coefficients of *S*(*z*) satisfy

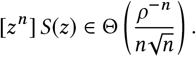

*Example.* Let 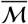 be restricted to the single motif (▭▭•), a special case of the local obstructions in the *Nussinov* Model described in [22]. Then, the O.G.F. of the set 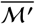 with *θ* = 1 is 1 + *z*^5^*T*^2^(*z*) and

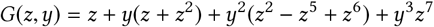

Next, we compute the resultant of the polynomial *P*(*z*, *y*) and its partial derivation on *y*

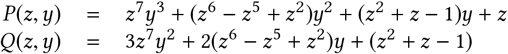

A numerical resolution of system locates the dominant singularity at *ρ* = 0.3834. We conclude that

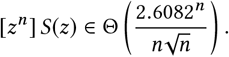

## 6 RESULTS

We implemented Algorithm 1, and the numerical procedure to compute the dominant singularity described in Section 5, in `Python3` using the pandas library and `SymPy` [35], a Python library for symbolic computing. Our implementation is available at: http://www.lix.polytechnique.fr/~ponty/?page=countingdesigns

### 6.1 Recovering the total number of secondary structures (*θ* = 3)

As a first test, we ran Algorithm 1, using the suboptimal defect 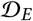 as our objective and no tolerance for suboptimality (*ε* := 0), based on `RNAfold` [30] (version 2.4.12 with default parameters), in order to detect local obstructions of small sizes (*k* ∈ [2, 4]). Unsurprisingly, but still reassuringly, our implementation returned three local obstructions, (), (•), and (••), corresponding to the *θ* = 3 minimal distance enforced by `RNAfold`.

Such local obstructions lead to a generating function

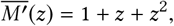

and our method produces the following asymptotic upper bound for the number of designable structures of size *n*:

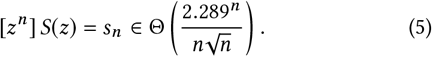

Those asymptotics match the ones reported by Hofacker *et al* [25].

### 6.2 Refined estimates for the phenotype space

We pushed our analysis further by using Algorithm 1 to compute the local obstructions of sizes up to 14 (hairpins) and up to 10 (internal loops and bulges). After removing the redundant motifs from the set, we manually computed the correcting terms *c*(*z*, *T*) to avoid double-counting structures compatible with several obstructions. Then, we applied the methodology of Section 5 to produce the dominant term of the asymptotics, along with the first-order estimates for the proportion of designable structures reported in Table 1.

**Table 1:**
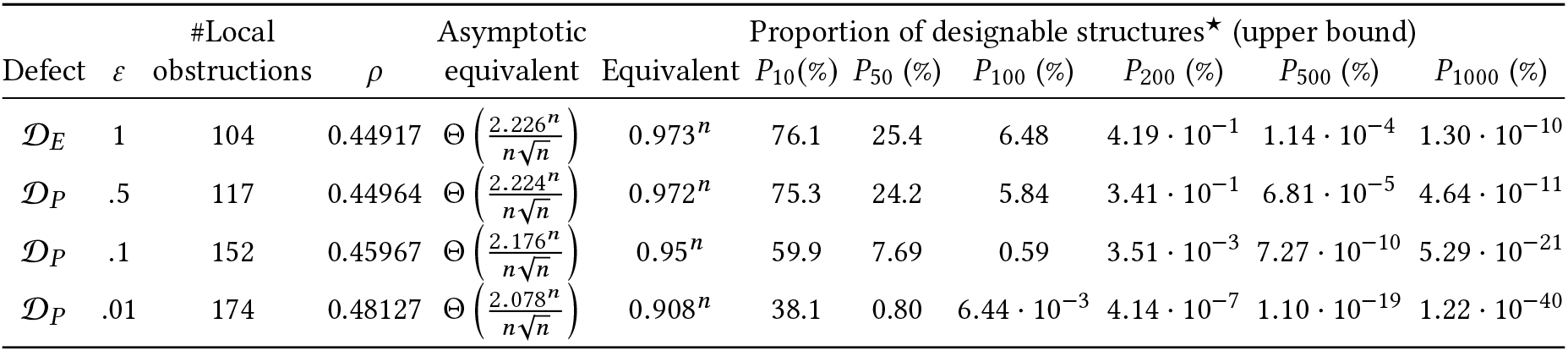
Collections of local obstructions of size up to 12, and their consequences on the proportion of actually designable secondary structures. ⭑ Proportions of designable sequences computed using an assumption of equal constants for the asymptotic leading terms of the number of secondary structures, respectively allowing and forbidding local obstructions.

#### 6.2.1 Inverse folding

In the classic setting of RNA design, the inverse folding, one attempts to design a sequence which admits a target structure as its unique MFE structure. This corresponds to choosing a suboptimal defect with *ε* = 1.

Our analysis reveal the existence of 104 motifs (after removal of redundant ones), an overwhelming majority of which contain isolated base pairs. Such motifs are expected, as they are heavily penalized, yet not explicitly forbidden (unless specified), by folding algorithms. Consecutive bulges, alternating on the 5’ and 3’ ends of an helix, also seem systematically suboptimal for the Turner model, a large interior loop being systematically favored as a candidate for the MFE. Finally, hairpin loops directly stemming from a multiloop are systematically discriminated, and a structure consisting of a larger unpaired stretch in the multiloop seem systematically favored.

Computing the dominant singularity yields *ρ* = 0.44917, which implies the following asymptotic upper bound on the number of secondary structures

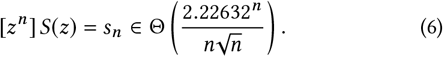

The probability for a secondary structure of size *n*, taken uniformly at random, to be designable is upper-bounded by *P*_*n*_ ∈ Θ(0.973^*n*^). Assuming the identity of constants involved in the leading terms of Equations (5) and (6), one concludes that, while about 3 4 of the structures of size 10 can be designed, this proportion quickly drops to less than 0.5% for RNAs of size 200, and reaches infinitesimal proportions (10^−10^%) for very large RNAs of size 1 000.

#### 6.2.2 Designing structures with large probabilities

Next, we analyze the probability defect 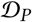, and investigate the impact of *ε* on the proportion of designable secondary structures. We consider 3 thresholds, *ε* ∈ {0.5, 0.1, 0.01}, associated with Boltzmann probabilities for the motif greater than 50%, 90% and 99% respectively. Executing Algorithm 1, followed by a removal of redundant motifs, led to the identification 117, 152 and 174 local obstructions respectively.

Interestingly, the *ε* = 50% case induces a dominant singularity of 0.44964, leading to a slightly slower asymptotic growth

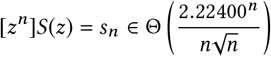

than for classic inverse folding. This is not entirely unexpected, since our definition of a valid design requires the target structure to be the sole MFE for the sequence. Thus, secondary structures satisfying some probability defect condition must also be solutions to the inverse folding problem. However, the observed divergence of the two singularities suggests that an exponentially small proportion (albeit with growth factor very close to 1) of MFE designs have Boltzmann probability greater than 50%.

For defect thresholds of 0.1 and 0.01 on the probability, the departure from the MFE design is much more pronounced, with respective singularities at 0.45967 and 0.48127 respectively, leading to asymptotic equivalents in

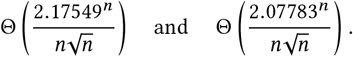

Again, assuming the equality of constants, we obtain proportions of designable structures bounded by *P*_*n*_ = 0.95^*n*^ and *P*_*n*_ = 0.908^*n*^ respectively. Those estimates support the notion of an extreme sparsity of designable structures in the folding space, with only three out of 10^−5^ (resp. 4 out of 10^−9^) structures being designable for *ε* = 0.1 (resp. *ε* = 0.01). These abysmal proportions are consistent with the popular belief, which rigorously holds for the homopolymer model [12], that the Boltzmann probability of the MFE structure decreases exponentially with the sequence length in a random, uniformly distributed, RNA sequence.

## 7 CONCLUSION

In this work, we have addressed the designability of RNA structures for a variety of design paradigms, thresholds and energy models. We have described a procedure for computing a list of local motifs whose presence represents an obstruction to the design task. This procedure is largely agnostic to the exact objectives of design, and holds for any design under mild assumptions (monotonicity of defects over loops). Using enumerative and analytic combinatorics techniques, we were able to automate the computation of asymptotic upper-bounds, revealing an overall sparsity of designable structures within the full conformational space, both in simple base-pair based and the complete Turner energy model. The number of designable structures still increases with the length of considered targets, but at a much slower rate than initially anticipated.

This work sets the stage for further analyses of designable structures, and unlocks a systematic way to address many further questions. For instance, the popular ensemble defect [48], could benefit from a more refined treatment using bivariate generating functions. Indeed, the ensemble defect is defined as an expectation, and is therefore fully additive on the Turner loops of the target secondary structures. One could therefore determine, through a trivial modificatioonf Equation (4), the bivariate generating function 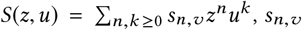 being an upper bound for the number of structures of size *n* having *v* ensemble defect. An application of the famous Drmota theorem [11] would then very likely provide sharper estimates, accounting for the accumulation of local defects rather than restricting to the worst one.

Enumerative aspects of this work could also easily be extended to secondary structures including algebraic types of pseudoknots. Indeed, multiple grammars have been shown to capture major pseu-doknot classes while, at the same time, allowing for a characterization of generating functions [36]. An enumeration of designable structures would greatly help in the parametrization of free-energy models, a key aspect of pseudoknot prediction programs which has so far greatly hindered the development of predictive methods [33].

Regarding the complexity of our method for building local obstructions, we strongly believe its exponential nature may be intrinsic to the problem. More precisely, we believe that the list of local obstructions may generically grow exponentially with the length of investigated motifs. Indeed, since structures can be seen as motifs in our definitions, the existence of a polynomially-bounded list of local obstructions would imply a polynomial-time algorithm for decided whether an RNA can be designed. Unfortunately, the problem was recently been shown to be NP-hard [4], appearing to rule out a polynomial-time alternative to Algorithm 1.

On a more positive note, Algorithm 1 can easily be modified to keep the list of suitable candidate sequences for each and every designable motif. This allows to greatly restrict the search space of classic design algorithms, but also suggests a promising strategy for hard design instances. As an illustration, while investigating our database of local obstructions, we discovered that lonely base pairs appear in a few designable motifs, usually considered unstable in the *Turner* model and difficult to design for. For example, the structure (((.(….).))) is the MFE structure of the RNA sequence UCAGCUUAUGGUGA. We also found that the motif ((‥(*)‥)) could be designable for some collection of sequences. Combining sequences adopting these two motifs, we could verify that an RNA sequence

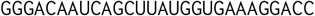

is predicted by `RNAfold` to adopt its unique MFE structure of

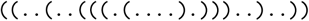

featuring two isolated base pairs, and a free-energy of −6.4kcal.mol^−1^, a stability unmatched across several runs of `RNAinverse` [24] and Nupack [48]. While this observation remains anecdotal, it is supported by the success of recent approaches using (partial) libraries of local motifs [3].

## A PROOF OF PROPOSITION 1

### PROPOSITION.

*For any defect* 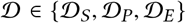, *sequence w*, |*w*| = *n, and structure* 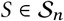, *one has*

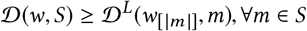

*where w*_[|*m*|]_ *is the restriction of w to the positions in m.*

PROOF. Let constraint *C* be the set of all (un)paired positions of *S \ m* plus the paired leaves and the root of *m* and 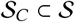 denotes the set of secondary structures of size *n* that are compatible with the constraint *C*. Then, we have the follow inequality

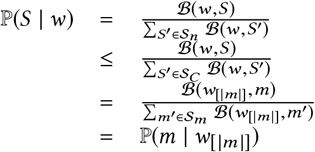

Therefore, 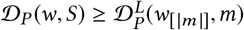.

Similarly,

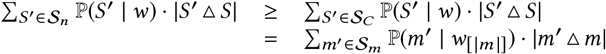

Thus, the inequality for 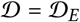

For the case 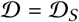, we consider *m*′ such that,

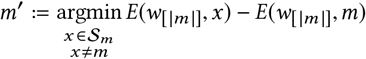

A such *m*′ exists since the set 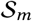 is finite. Let 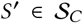 be the secondary structure containing *m*′ at the position of *m*. We have,

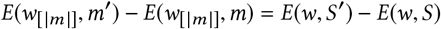

In addition, we have, by definition, 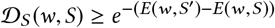, which implies the inequality 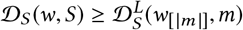

## B PROOF OF PROPOSITION 2

### PROPOSITION.

*For every defect* 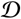*and energy model considered in this work, one has* 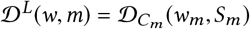.

PROOF. The schema of proof is similar to the above one. Let 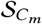 be the set of secondary structures of length |*S*_*m*_| that are compatible with the folding constraint *C*_*m*_. One can observe that the way to make the completion structure is a bijection from 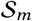 to 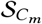. Let *m*′ be a motif equivalent to *m* and 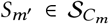 be its minimum completion structure. The energy of the structure *S*_*m*_(resp. *S*_*m*_′) is the sum of the energy contribution with the motif *m* (resp. *m*′) and with the constrained part, which is the same for both structures. The later one is a constant for a given sequence. For the reason of simplicity, we denote it by *E*_*C*_. Then, for a given sequence *w* and its minimum completion *w*_*m*_, we have

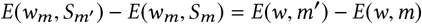

This proves the proposition for the case where 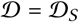

For the case of the Probability Defect 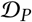,

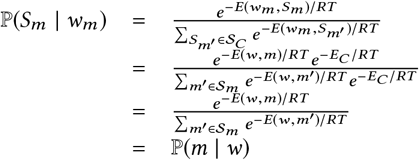

Thus, the equality.

Furthermore, for any motif 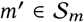 and its minimal comple-tion *S*_*m*′_, the base pair distance are equal between motifs *m*′ and *m* and between their minimal completion structures, |*m*′ Δ *m*| = |*S*_*m′*_ Δ *S*_*m*_|, because the completion part is same for both motifs. Therefore, the equality for the case of the ensemble defect 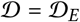.

